# Patient-derived monoclonal anti-HPA-1a can induce platelet activation through FcγRIIa

**DOI:** 10.64898/2026.09.10.750554

**Authors:** Janita J. Oosterhoff, Suze R van Brummelen, Marco Giulini, Iris M De Cuyper, Remco Visser, Rick Kapur, Christoph Gstoettner, Elena Domínguez Vega, Thijs de Vos, Suzanne Hofstede-van Egmond, Mark S Cragg, Manfred Wuhrer, Leendert Porcelijn, Masja de Haas, Alexandre MJJ Bonvin, C. Ellen van der Schoot, Gestur Vidarsson

**Author notes:** Corresponding author: Gestur Vidarsson.

## Abstract

Fetal and neonatal alloimmune thrombocytopenia (FNAIT) is a pregnancy-associated disorder caused by maternal alloantibodies targeting paternally inherited human platelet antigens (HPAs). These antibodies traverse the placenta and bind to fetal platelets, causing thrombocytopenia and potentially severe complications including intracranial hemorrhage (ICH). Beyond platelets, antibodies may also target other fetal cells such as endothelial cells and placental trophoblasts thereby exacerbating disease severity. Anti-HPA-1a antibodies, the primary cause of FNAIT, display functional heterogeneity due to differences in epitope specificity, glycosylation, which complicate disease severity prediction. Here, we report the structural and functional characterization of two novel anti-HPA-1a antibodies, D-204 and M-204, derived from a mother with a severe FNAIT case. In comparison to D-204 and other existing anti-HPA-1a antibodies such as B2G1 and 26.4, M-204 bound αIIbβ3 with different kinetics, reaching only half-maximal binding, as determined by flow cytometry. Notably, only M-204 induced platelet aggregation. To investigate the underlying mechanism behind this aggregation, various antibody formats were generated (Fab, Fab2, bispecific, and IgG-Fc dead). Aggregation required both Fab-arms to engage HPA-1a and intact binding to FcγRIIa. Structural models and crystal structures suggest the dual engagement of both Fab-arms is required for FcγRIIA-engagement which is a unique feature of M-204 among known HPA-1 monoclonals. Our findings highlight the functional diversity of anti-HPA-1a antibodies and provide new insights into their pathogenic mechanisms.

## Introduction

Fetal and neonatal alloimmune thrombocytopenia (FNAIT) is a pregnancy-associated disorder that can result in intracranial hemorrhage (ICH) and organ bleeding during pregnancy or in the neonatal period ^1,2^. This condition occurs when maternal alloantibodies, generated in response to paternally inherited human platelet antigens (HPAs), traverse the placenta via active transport^1,3^. Once in the fetal circulation, these antibodies bind fetal platelets thereby causing thrombocytopenia^4^. Beyond platelets, these antibodies can also target other fetal cells expressing the involved antigen including endothelial cells and placental trophoblasts, amplifying disease symptoms^5,6^. The clinical spectrum ranges from asymptomatic to severe complications. Timely treatment with intravenous immunoglobulins (IVIG), and potentially with future FcRn inhibitors, can prevent severe disease^7^. However, the absence of laboratory tests to identify the relatively small fraction of alloimmunized pregnancies at high risk for antenatal ICH, hampers the implementation of routine screening for the presence of anti-HPA alloantibodies. Therefore, at present antenatal treatment is only given to women with a history of FNAIT ^8–10^.

FNAIT is most commonly caused by antibodies directed against the HPA-1a antigen which occurs as a result of a polymorphism found in the β3 integrin subunit^11^. The classical view of FNAIT proposes fetal platelets to be responsible for immunization which express high levels of the fibrinogen receptor, integrin αIIbβ3^12^. However, trophoblast cells of the placenta may also act as an important immunogenic source due to expression of the vitronectin receptor, αvβ3^13^. These cells invade maternal uterine tissue and are in contact with maternal stroma and immune cells^5,6^. The clinical heterogeneity of FNAIT might be reflected in the specificity and functional properties of anti-HPA-1a antibodies^14^. Some anti-HPA-1a antibodies directly impair cellular processes such as platelet adhesion to fibrinogen^15,16^, endothelial cell survival^17–19^ and trophoblast invasion^5,6^. Others exert direct effects by blocking integrin function and modulating integrin activation status^20^. Differences in IgG-Fc-glycosylation contribute to variations in their capacity to activate complement and recruit effector cells^21–24^. It has been suggested that anti-HPA-1a antibodies exhibit distinct preferences for their epitope with some targeting αIIbβ3 or αvβ3 selectively and others binding both^17,25^. However, more recent findings contradict these claims, as anti-HPA-1a sera depleted using either αIIbβ3-or αvβ3-expressing cells, leave no reactivities to the reciprocal counterpart. Similarly, eluted antibodies from such cells also show no preference for either antigen. In addition, in a more recent analysis, the glycosylation of such antibody pools differ, in particular with respect to afucosylation, the unique fingerprint of which also strongly suggest that anti-HPA-1a do not tend to discriminate between αIIbβ3 or αvβ3^26,27^.

Recently, we reported the isolation of two novel anti-HPA-1a antibodies, clone D-204 and M-204, from a mother with a severe FNAIT case^28^, and solved their structures^29^. In the current study, we characterized these antibodies in terms of epitope specificity and functional effects using various target cells including HEK-293F cells transduced with integrin αIIbβ3 or αvβ3, HUVECs or platelets.

## Materials and methods

### Generation of antibody formats

Variable heavy (VH) and light (VL) chain sequences of anti-HPA-1a antibodies B2G1, 26.4, D-204 and M-204 have been described previously^28,30,31^. M-204 without a Fab-glycan was made by mutating the N-glycosylation site in FR3 back to germline (N72D). Monoclonal antibodies were cloned and produced as IgG1 (IGHG1*03) in HEK-293F cells as described before^28,32,33^. For Fab production, a shortened IgG constant heavy was generated ending with a specific site above the hinge (KSCDKT). Fc dead variants were generated by mutating specific residues in the IgG-Fc namely P329G, L234A, L235A (PGLALA), N297A and H435A which render the Fc part inactive for binding to FcγR and complement. Human embryonic kidney 293F (HEK-293F) cells were maintained in Freestyle™ Expression Medium (Thermo Scientific) under controlled conditions of 37°C and 5% CO₂ with agitation at 125rpm. Supernatants were collected 6 days post-transfection and subjected to purification using a 5-mL protein A or G column (GE Healthcare), for full antibodies or Fc dead variants respectively. Fab fragments were purified using CaptureSelect CH1-XL (Thermo Scientific). The purified eluates were subsequently dialyzed against 5 mM sodium acetate (pH4.5) and stored at −20°C for downstream applications. F(ab’)2 fragments were produced by cleaving full antibodies overnight with pepsin (1:100 pepsin:IgG, based on mass) in 0.1M sodium citrate at 37C°. Cleavage was stopped by raising the pH with 1M Tris pH8.0. Samples were purified using CaptureSelect CH1-XL and dialyzed against sodium acetate. For duobody generation, Fab-arms of F405L anti-HPA-1a and K409R anti-TNP, or when indicated, without an irrelevant Fab (single anti-HPA-1a Fab), were exchanged and verified as described before^34,35^. Anti-HPA-1a antibodies were labeled with DyLight^TM^ 650 NHS Ester (Thermo Scientific) according manufacturer’s instructions.

### SDS-PAGE

Antibodies and antibody fragments were tested in SDS-PAGE under reducing and non-reducing conditions. Samples were dissolved in NuPage LDS Sample Buffer (Thermo Scientific) and incubated for 5min at 95C° or 70C° in the presence of 2-mercaptoethanol (Sigma-Aldrich) or 20mM iodoacetamide (Sigma-Aldrich) for reduced and nonreduced conditions, respectively. Separation of samples was performed using a NuPage^TM^ 12% Bis-Tris protein gel (Invitrogen) in MOPS SDS running buffer. After staining with InstantBlue (Abcam) for 30min and destaining in distilled water, the gel was visualized using Gel Doc (Universal Hood III, Bio-Rad).

### NanoLC-MS

To generate antibody subunits, antibodies were diluted in phosphate-buffered saline (PBS) to a concentration of 1μg/μL. Subsequently, the antibodies were digested below the hinge region using IdeS protease (Genovis) at a ratio of 2U/μg antibody. After incubation at 37°C for 2h, the Fab and Fc/2 subunits were denatured and reduced for 30min at room temperature using Guanidine hydrochloride and TCEP at a final concentration of 4M and 20mM, respectively. The reduced subunits were analyzed on an U3000 nanoLC system (Thermo Fisher Scientific). For each analysis, 1μL of sample were injected onto a C4 trap column (5.0 × 0.3mm i.d., Acclaim™ PepMap™, 300Å pore size; Thermo Fisher Scientific) using a flow rate of 15μL/min. Trapping of the samples was performed using isocratic conditions (H2O + 0.1% trifluoroacetic acid; TFA; Merck), at a temperature of 60°C for 5min. The subunits were separated using a diphenyl reversed phase column (150.0 × 0.1mm i.d, Halo® Bioclass, 1000Å pore size; Advanced Material Technology) at a flow rate of 1.0 μL/min. The separation column was placed in a butterfly heater (Phoenix S&T) set to 80°C. A multi-step gradient using as mobile phase A H2O (ELGA Labwater) + 0.1% TFA) and mobile phase B (acetonitrile (Actu-All Chemicals, Oss, the Netherlands) + 0.1% TFA) was programmed as follows: 20.0% B for 5 min, 20.0%–32% in 1min, 32–55.0% B in 12min, 55.0–70.0% B in 2min, 90.0% B for 5min, and 20% B for 5min. The U3000 nanoLC system was hyphenated to a QTOF mass spectrometer (Maxis HD, Bruker) via a nano-ESI source (CaptiveSpray, Bruker). This source allows the usage of an acetonitrile-enriched nitrogen gas at a pressure of 0.1 (nanoBooster; Bruker). The MS parameters were set as following: Dry gas flowrate was set to 3L/min, dry gas temperature to 220°C. The MS was operated in mass range of m/z between 600 to 6000 in positive ionization mode with a capillary spray voltage of 1050V. To achieve efficient declustering of the molecules an in-source CID of 80eV, a collision cell RF of 2000Vpp, a quadrupole ion energy of 5eV and collisions cell energy of 7eV were used. The pre-pulse storage was set to 20μs, and the transfer time to 150μs.

### Cell culture

HEK-293F cells stably transduced with either integrin αIIbβ3 or αvβ3 bearing the HPA-1a epitope were generated as described before ^26^. In short, αv and HLA-class I were knocked out prior to transducing the cells with integrin constructs. Cells were maintained in Freestyle™ Expression Medium (Thermo Scientific), supplemented with 5µg/ml blasticidin, 100U/ml penicillin and 100µg/ml streptomycin (Thermo Scientific), under controlled conditions of 37°C and 5% CO₂ with agitation at 125rpm. HUVECs (pools of 5 donors, Promocell) were cultured in Endothelial Cell Growth medium 2 (EGM-2, Promocell) supplemented with 5% FCS, 4mM L-glutamine (Thermo Scientific), 100U/ml penicillin, and 100µg/ml streptomycin (Thermo Scientific) at 37°C with 5% CO2 in Nunclon Delta Surface flasks (Thermo Scientific) coated with 50µg/ml rat tail collagen I (Thermo Scientific). HUVECs were washed with PBS and detached with trypsin/EDTA (0.05%/0.02% m/v) solution. Trypsin was neutralized with Trypsin Neutralizing Solution (Lonza) before cells were resuspended in fresh EGM-2.

### Flow cytometry

To determine anti-HPA-1a antibody opsonization, anti-HPA-1a antibodies were titrated in PBS/0.5%BSA and allowed to bind antigen on HEK-293F cells expressing integrin αIIbβ3 or αvβ3 (0.5×10^5^ cells/ml), HUVECs (0.5×10^5^ cells/ml) or platelets (0.5×10^6^ platelets/ml) for 30min at 4°C (room temperature for platelets). After three washes, the cells were incubated with either FITC-or PE-conjugated mouse anti-human IgG (JDC-10, Southern Biotech) for 30min at 4°C or room temperature. For testing more secondary reagents, clone G18-145 (MaH-IgG-V450, BD Biosciences), clone SB81a (MaH-kappa-PE), and anti-IgG-CH1 (nanobody, Thermo Scientific) were used, the latter in combination with Strep-APC (Biolegend). Cells were washed twice and measured on a FACS LSRFortessa^TM^ or BD FACS Fortessa (BD Biosciences). For absolute platelet counting by FACS, CountBright Absolute Counting Beads (Thermo Scientific) were used according to manufacturer’s instructions. Fixation before staining was performed by incubation of platelets with 1% paraformaldehyde (PFA) for 10min at room temperature.

### Light transmission aggregometry

Platelets were isolated from citrated whole blood from healthy anonymous volunteers with known HPA-1-type and centrifuged at 150g for 20min at room temperature. The platelet-rich plasma was harvested and supplemented with 300nM prostaglandin E1 (PGE1, Sigma Aldrich) before centrifugation at 480g for 15min to pellet platelets. Platelets were resuspended in warm PIPES/saline/glucose (PSG) buffer (5mM PIPES, 145mM NaCl, 4mM KCl, 50µM Na2HPO4, 1mM MgCl2 and 5.5mM D-glucose, pH6.8) containing 300nM PGE1 followed by centrifugation at 480g for 15min. The washed platelets were resuspended in FACS buffer (PBS, 0.5% BSA), counted (Beckman Z1 counter) and diluted to 300×10^6^ platelets/ml. Platelet aggregation was performed in the Chrono-log 490 aggregometer at 37C° at 1200rpm. After 5min of preincubation, with or without AT10^36^ antibodies (50µl) to block FcγRIIa were added manually to 200µl of platelets and light transmission was recorded for 20min. Results were reported as the maximum amplitude of light transmission which was used as reflection of the aggregation potential.

### Modelling

AlphaFold2-multimer (AF2)^37^, AlphaFold3^38^, and Immunebuilder (IB)^39^ were used to generate predictions for the apo conformation of the M-204 and D-204 antibodies, while AF2, AF3, and HADDOCK^40,41^ were used to predict the structure of their complex with αIIbβ3 integrin proteins, based on the PDB 3FCS^42^ ^40^. Structural ensembles, known to be beneficial for the docking, were used in both cases combining AF2 and IB models^43,44^. The D-204 ensemble included eight models while the M-204 ensemble included only two. The docking was performed on local resources using HADDOCK3 workflow with increased sampling (5000 models) to compensate for the presence of an ensemble. Fraction of common contacts clustering was performed after the rigid-body docking stage and at the end of the workflow^45^. Docking restraints were generated by setting the surface exposed residues on the antibodies’ CDR loops as active, thus being penalized if not in contact with the antigen. On the antigen side, LEU33 was defined as active as well, with its neighboring (within 10A) surface-exposed amino-acids being labelled as passive and therefore not paying any restraint penalty if they were not in contact with the antibody.

### Data analysis and statistics

Flow cytometry data were analyzed with FlowJo software v10.8.1. Data were plotted and statistically analysed using Graphpad prism software (version 10). Specific tests are indicated in the figure legends. Mass spectra was deconvoluted using the DataAnalysis software from Bruker. NetNGlyc (https://services.healthtech.dtu.dk/services/NetNGlyc-1.0/) was used for prediction of N-glycosylation sites. Structural visualizations were performed with UCSF ChimeraX version 1.5.

## Results

### Structural characterization of anti-HPA-1a antibodies D-and M-204 reveals Fab glycosylation in M-204

To investigate the structural composition of anti-HPA-1a monoclonal antibodies D-204 and M-204, the antibodies were produced as full IgG and Fab fragment followed by characterization by sodium dodecyl sulfate polyacrylamide gel electrophoresis (SDS-PAGE) (Fig. 1A-B). Under non-reducing conditions, a band around 150kDa or 50kDa was found for the full antibodies and Fab fragments, respectively (Fig. 1A). Interestingly, the Fab fragment of M-204 was of higher molecular weight which was also observed under reduced conditions (Fig1B). Potential N-glycosylation sites were identified within the variable region in the FR3 of M-204 (NTS), but also in the CDR2 of D-204 (NGS).^28^ The actual glycosylation status was then verified by mass spectrometry (MS), where the antibodies were subjected to hinge-region digestion, reduction and subsequent analysis of the resulting subunits (LC, Fc/2 and Fd’) by nanoLC-MS (Fig. 1C). For D-204, the predominant signal corresponded to the Fd’ subunit without Fab glycosylation, exhibiting a mass of 25564Da. Only trace amounts of Fab glycosylation at 27500 Da were detected, although these minor signals were not clearly distinguishable in MS due to their low intensity, but consistently supported by a faint band observed in the gel (Fig. 1C). In contrast, analysis of M-204 indicated that nearly all Fd’ subunits were glycosylated, represented by a distribution of glycoforms in the deconvoluted mass spectrum at higher masses with the primary glycosylated peak having a mass of 27442Da while a minor 25340Da non-glycosylated Fd’ subunit peak was also present. Thes 27442Da mass and deducted composition is compatible with both a tri-antennary glycan or a bisected biantennary glycan based solely on their intact mass. The latter was previously found to predominate in Fab glycosylated antibodies and therefore peaks were annotated assuming bisection rather than tri-antennarity^46^.Mutation of the N-glycosylation consensus sequence (NTS) in M-204 resulted in the detection of only non-glycosylated Fd’ subunits by both SDS-PAGE and nanoLC-MS (Fig 1A-C).

**Figure 1.**
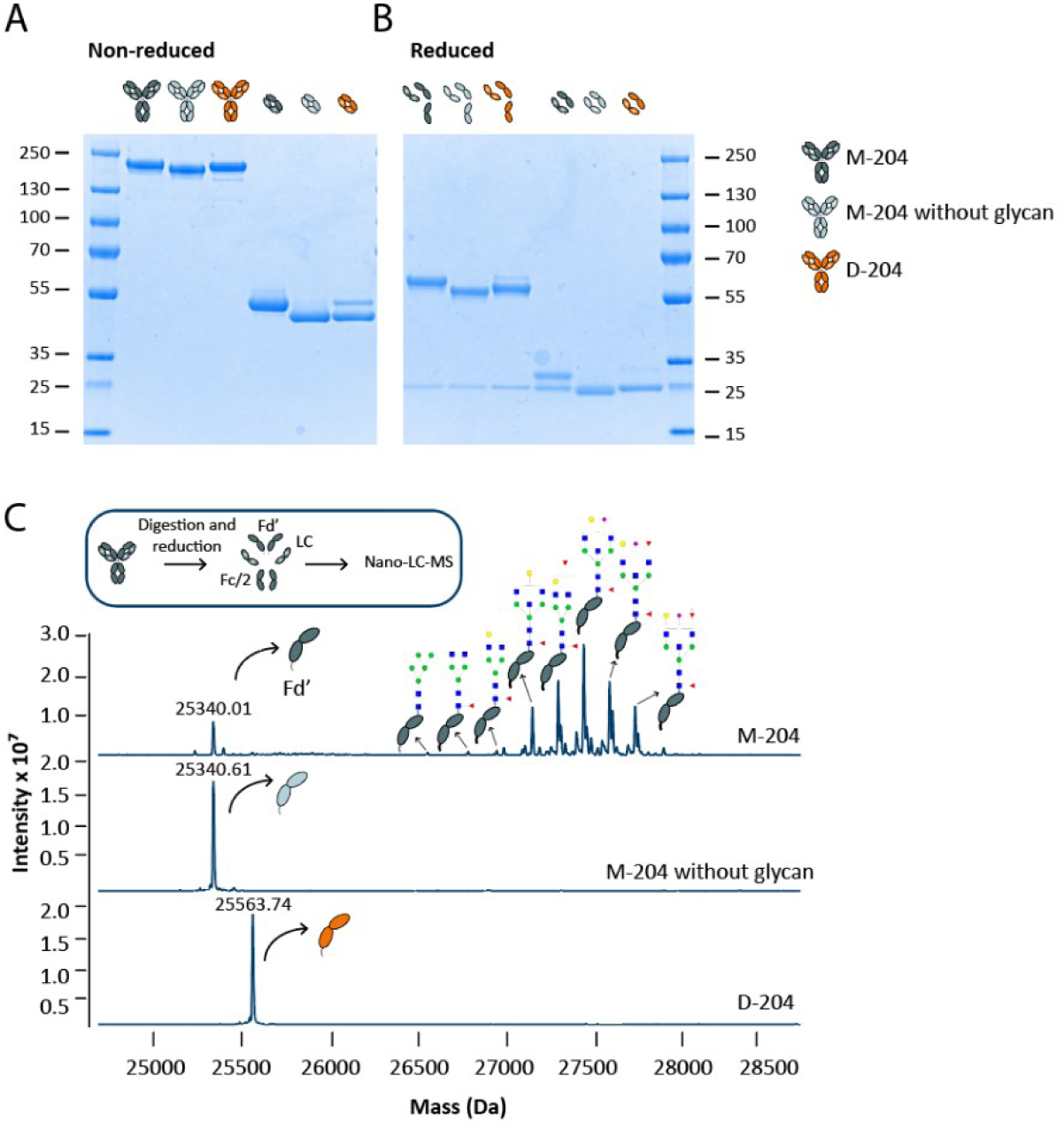
SDS-PAGE and nanoLC-MS analysis of D-and M-204 confirm the presence of a Fab glycan exclusively in M-204. **(A)** SDS-PAGE of non-reduced samples shows bands at approximately 150kDa for full IgG and 50kDa for Fab fragments. **(B)** SDS-PAGE of reduced samples reveals bands at 50/25kDa and 25kDa for full IgG and Fab fragments, respectively. The bands corresponding to M-204 are positioned higher compared to D-204, a difference reversed in M-204 without glycan where the IgG-Fc glycosylation site (NTS) was mutated (DTS). **(C)** Nano-LC-MS of the Fd’ subunits confirms the presence of a glycan in the Fab fragment of M-204 showing clear signs of Fab glycosylation.

### M-204 reaches halfway saturation upon binding to αIIbβ3-expressing platelets coinciding with reduced platelet count

To assess the binding dynamics of anti-HPA-1a antibodies D-204 and M-204, as well as the previously described antibodies 26.4 and B2G1, titrations were performed on HEK-293F cells stably transduced with either integrin αIIbβ3 or αvβ3 (Fig. 2A, C, Supplemental Fig. 1). M-204 exhibited differential kinetics for these two HPA-1a containing β3 antigens depending on the expressed α-chain, which was not observed for any of the other anti-HPA-1a monoclonals. When binding αIIbβ3-expressing cells, M-204 only reached approximately half-maximal saturation values which was not observed for αvβ3 (Fig. 2C). M-204 consistently exhibited a lowered plateau phase, although the extent varied depending on the secondary reagent used or the use of directly labeled monoclonals. (Supplemental Fig. 2A-E). The presence of the Fab glycan did not appear to affect the binding characteristics as M-204 without glycan behaved similarly compared to its glycosylated counterpart (Fig. 2B). To validate these results on cells expressing endogenous integrin αIIbβ3 or αvβ3, we titrated the antibodies on αIIbβ3-expressing platelets and HUVECs obtaining similar results (Fig. 2B, D). This signal reduction was not observed on fixed platelet (Supplemental Fig. 1F). When investigating the platelet population by flow cytometry, increased number of aggregates were observed in the presence of M-204, as indicated by elevated forward/side scatter (FSC/SSC) (Fig. 2E,F). Additionally, the reduction in platelet counts suggested platelet loss due to aggregate formation (Fig. 2G).

**Figure 2.**
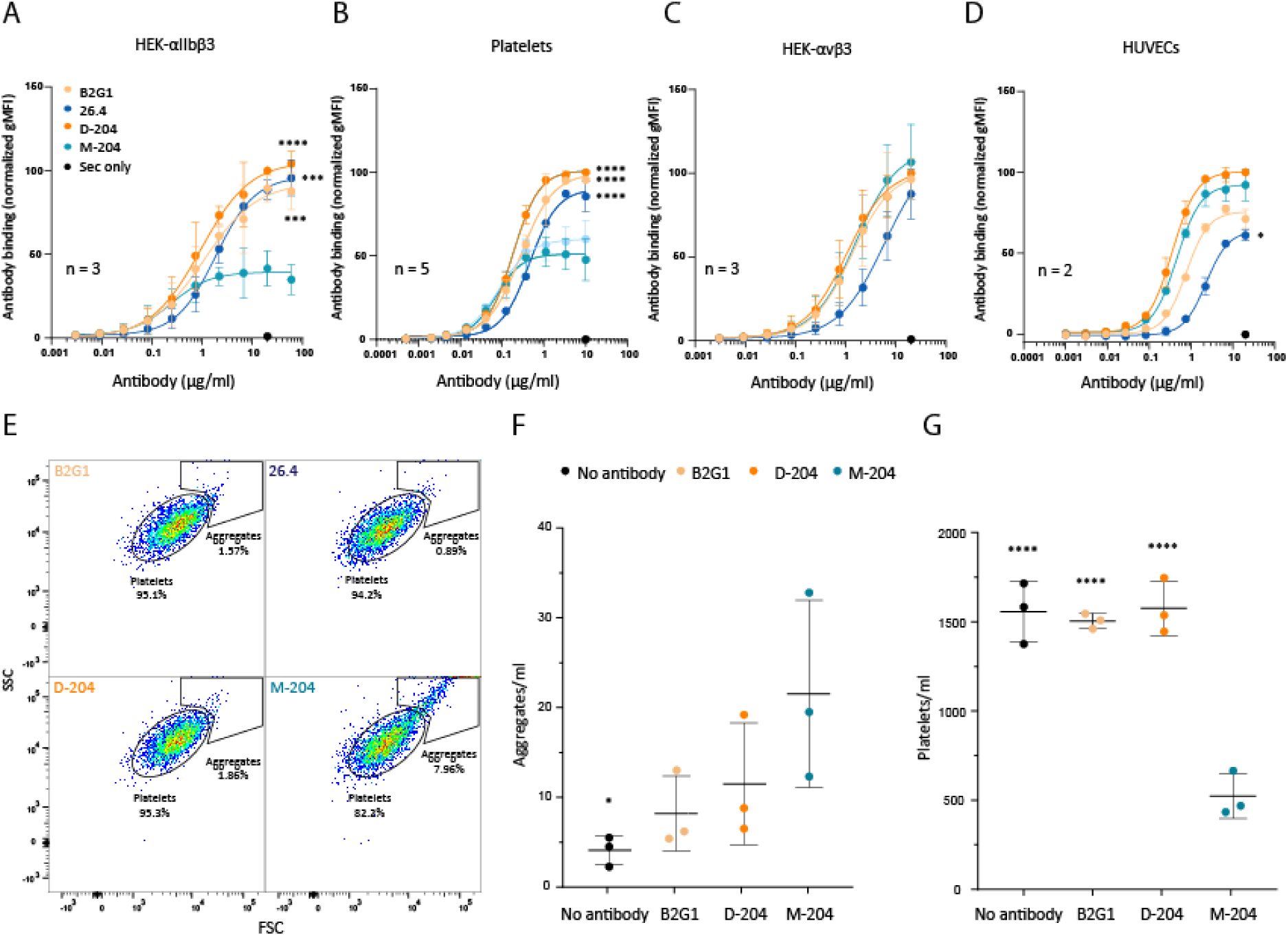
M-204 shows lower saturable binding to αIIbβ3, but not αvβ3, and induces platelet aggregation. **(A)** Antibody binding to HEK-293F cells expressing αIIbβ3, **(B)** platelets with the HPA-1a/1a genotype, **(C)** HEK-293F cells expressing αvβ3 and **(D)** HUVECs. **(E)** Representative dot-plots from platelets incubated with 10µg/ml monoclonal antibody. The platelet population as well as platelet aggregates are gated. **(F)** The aggregates and **(G)** platelet counts were quantified using counting beads. **(A-D)** Staining of cells was plotted as gMFI normalized towards 20µg/ml D-204. **(A,C)** Data is presented from three independent experiments or **(B)** five independent platelet (HPA-1a/1a) donors or **(D)** 2 independent experiments using flow cytometry. **(A-D, F-G)** Error bars indicate standard deviations, and statistical analysis was performed on the highest concentration using one-way ANOVA with Dunnet’s multiple comparison test comparing to M-204. Statistically significant differences are indicated by asterisks: *<0.1, ***<0.001, ****<0.0001.

### Aggregation of platelets by M-204 requires bivalent binding

We then performed light transmission aggregometry to specifically study platelet aggregation induced by M-204 in more detail (Fig. 3A). Aggregation was induced upon incubation with M-204 but not D-204, B2G1 or 26.4 (Fig. 3A-B). Platelet aggregation caused by M-204 was concentration-dependent with an optimal effect observed at 2 µg/ml (Fig. 3C-D). Curiously, aggregation decreased at higher antibody concentration, perhaps suggesting that bivalent antibody binding, where both Fab arms engage antigen, is required for this effect. At super-saturating concentrations, antibodies may predominantly bind monovalently, occupying only one Fab arm per antigen. In contrast, at lower concentrations, sufficient antigen density and lateral mobility may allow for bivalent binding^47^. However, other elements must also be involved as the aggregation potency was less sensitive to antibody concentration for heterozygous HPA-1a/1b platelets. This is highly relevant in FNAIT where the fetal genotype is always heterozygous in naturally induced pregnancies.

**Figure 3.**
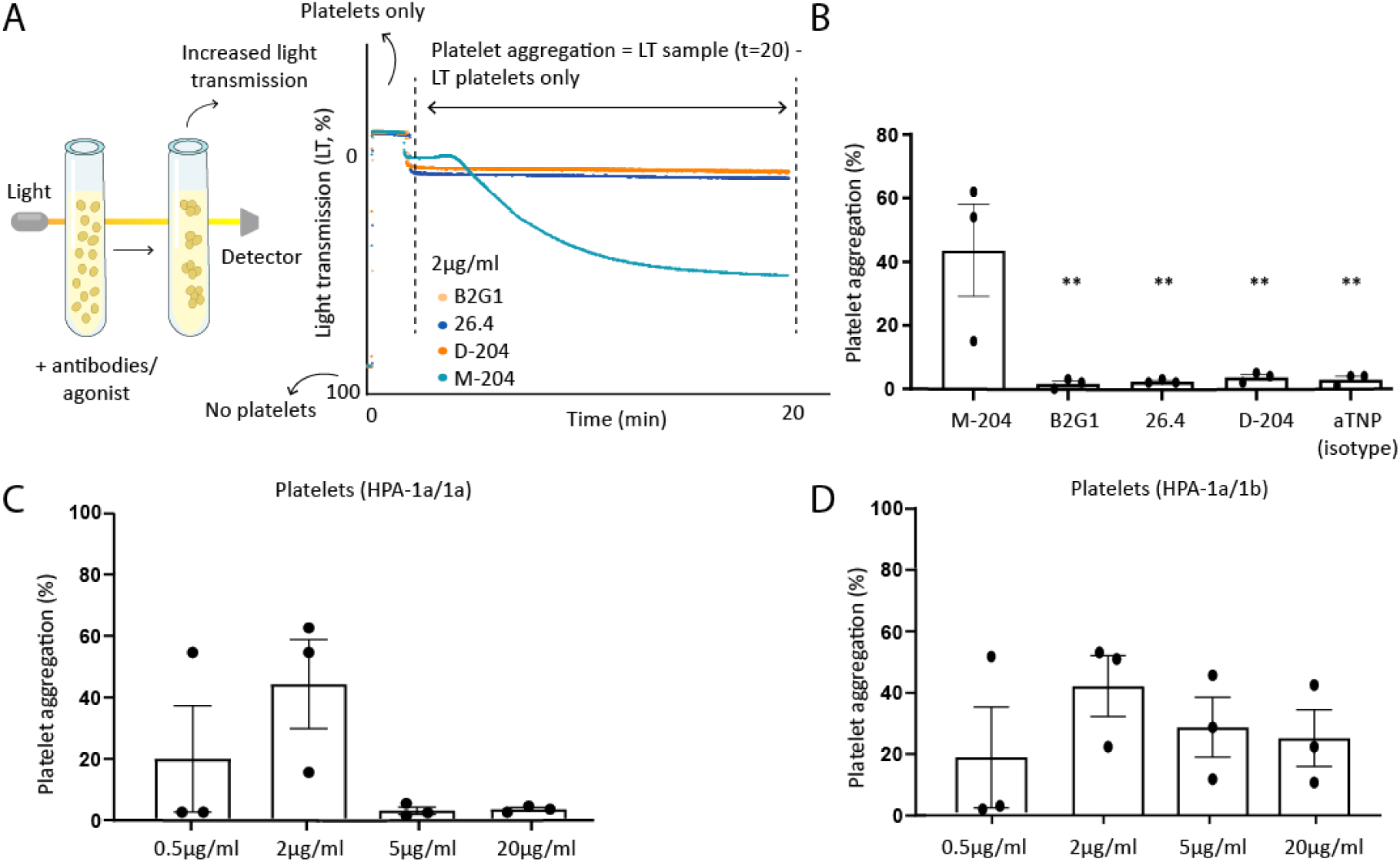
M-204 causes platelet aggregation which is concentration-dependent. **(A)** Schematic representation of light transmission aggregometry (left) and (right) a representative example of different monoclonal antibodies added to HPA-1a/1a platelets and the effect on light transmission (%) which represents aggregation. Light transmission was measured for 20min upon addition of antibodies. The maximum amplitude was used as readout after subtraction of background signal of platelets only. Medium only without platelets was used as positive control for 100% light transmission. **(B)** Platelet aggregation induced by anti-HPA-1a monoclonal antibodies M-204, B2G1, 26.4, D-204 and aTNP isotype as negative control, tested on HPA-1a/1a platelets. **(C)** M-204 was titrated on platelets with the HPA-1a/1a and **(D)** HPA-1a/1b genotype. **(B-D)** Data is presented from three different donors. Error bars indicate standard deviations and statistical analysis was performed using paired ANOVA with Dunnet’s multiple comparison test comparing to M-204 or 2µg/ml. Statistically significant differences are indicated by asterisks: **<0.01.

### M-204-induced platelet aggregation is FcɣRIIa-mediated

We then investigated the requirement for both Fab arms to induce platelet aggregation. First, Fab fragments of the HPA-1a monoclonals were generated and titrated for binding to HPA-1a/1a platelets. Unlike the full-length M-204 IgG (Fig. 2C), the M-204 Fab fragment reached the highest plateau phase compared to the other monoclonals (Fig. 4A), yet failed to induce platelet aggregation (Fig. 4B). This raises the question whether the lack of an Fc tail or the inability to bind bivalently and crosslink integrins was responsible for the absence of platelet aggregation. We therefore tested F(ab’)2 fragments and a Fc dead variant (incorporating PG-LALA, N297A and H435A mutations preventing binding to FcɣRs and FcRn)^48^ neither of which induced aggregation (Fig. 4B, Supplemental Fig. 3). Blocking FcɣRIIa on platelets with AT10 antibody was sufficient to prevented M-204-induced platelet aggregation, suggesting that platelet-FcɣRIIa is required for inducing their aggregation (Fig. 4B). To further dissect the underlying mechanism, we generated a bispecific (duobody platform^49^) targeting HPA-1a with one arm and TNP with the other arm. At high concentration the M-204 bispecific showed increased maximal binding overcoming the half-maximal binding seen by the other monoclonal antibodies (Fig. 4C), suggesting that intact M-204 to preferentially binds the HPA-1a epitope with both Fab arms. The bispecific antibody completely lost the ability to induce platelet aggregation, indicating that both bivalent binding and FcɣRIIa engagement are required for M-204-mediated platelet aggregation. This unique behavior of M-204 was also independent of it’s Fab-glycan as the aglycosylated (DTS) mutant behaved similarly to wildtype M-204 (Fig. 4D).

**Figure 4.**
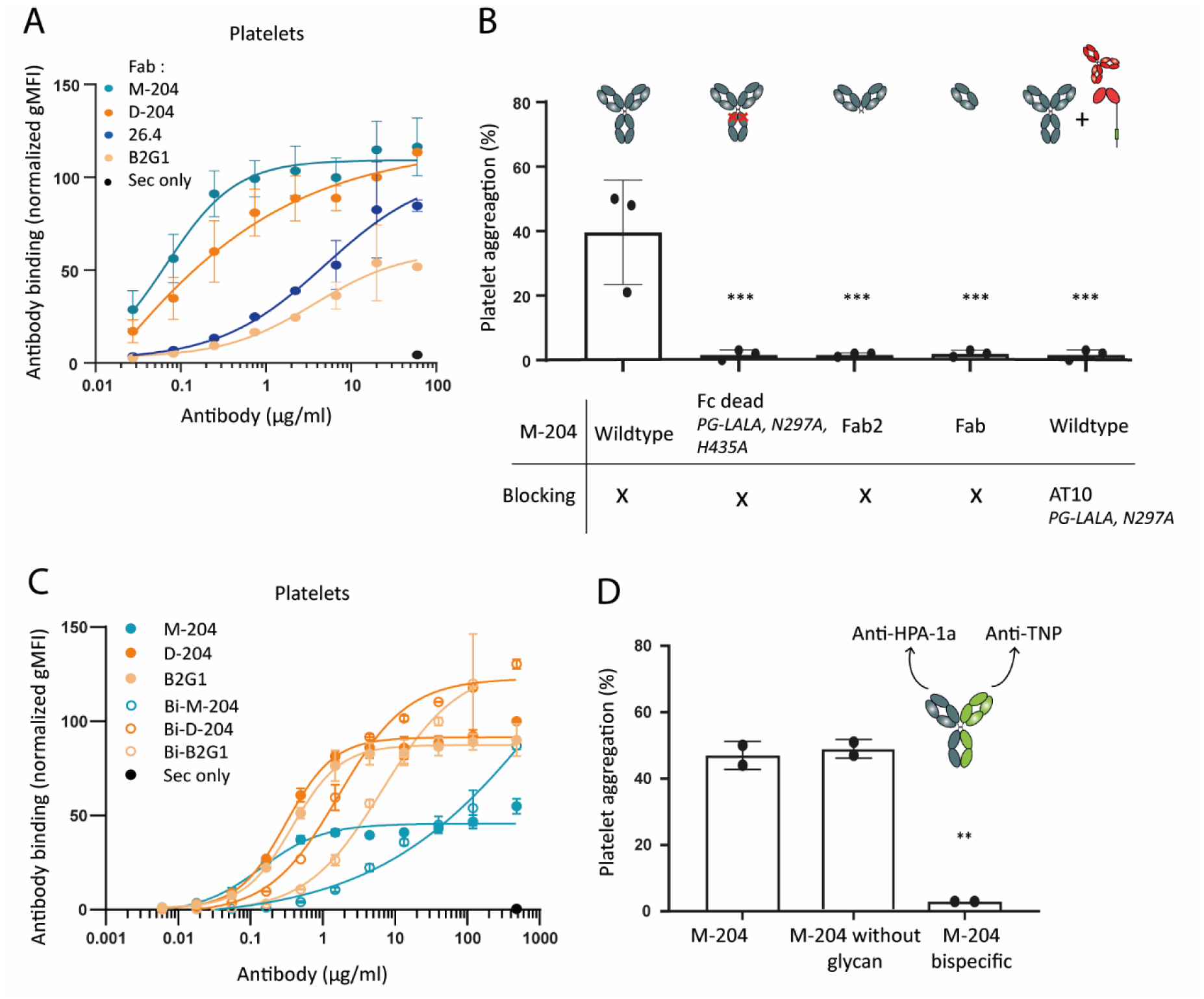
Both Fab arms and the Fc domain of M-204 are required to induce platelet aggregation. **(A)** Titration of anti-HPA-1a Fab fragments on HPA-1a/1a platelets. Anti-CH1 and Strep-APC were used as secondary detection reagents. **(B)** Several variants of M-204 were generated including an Fc dead variant with PG-LALA, N297A and H435A mutations, Fab2, and Fab fragments, all tested at 2µg/ml. The AT10 antibody with PG-LALA and N297A mutated backbone, targeting FcɣRIIa, was used to block aggregation of 2µg/ml M-204 full antibody. **(C)** Titration as in Fig. 1b of the indicated monoclonal or bispecific duobodies with single-Fab arm of the indicated HPA-1a clones on HPA-1a/1a platelets. **(D)** M-204 without glycan and M-204 bispecific (Duobody with an irrelevant anti-TNP specificity for the second Fab) were tested for their capacity to induce platelet aggregation. **(A-B)** Data is presented from three or **(C-D)** two different donors. Error bars indicate standard deviations and statistical analysis was performed using one-way ANOVA with Dunnet’s multiple comparison test comparing to M-204 at **(A)** 20µg/ml. Statistically significant differences are indicated by asterisks: **<0.01.

### Alternative binding modes of D-204 and M-204

To gain further understanding of why M-204 and D-204 have altered propensity to activate platelets, we attempted to test how these antibodies bind to the β3-containing HPA-1a epitope. First, the Fabs were modelled using AlphaFold2-Multimer (AF2^37^), AlphaFold3 (AF3^38^), and ImmuneBuilder (IB^39^). The obtained models showed high values of confidence (loop pLDDT always higher than 80) and suggested a fundamental difference between the two antibodies, with M-204 containing an extended CDR1 loop in the light chain (Supplemental Fig. 4A), while D-204 showed a relatively flat CDR-interface (Supplemental Fig. 4B). Modelling of these Fabs in complex with αIIβ3 integrin was first attempted with AF2 and AF3, resulting in predictions with poor confidence metrics (AF2 ipTM < 0.3, AF3 ipTM < 0.6) and few or no contacts between the antibody chains and the epitope amino acid LEU33. Therefore, we used HADDOCK^41^ to dock both M-204 and D-204 clones with the available αIIβ3 and αVβ3 integrins (PDB-ID 3FCS) and inspected the top ten clusters of each run (Supplemental Fig. 4E). During this work, crystal structures of the complexes between M-204 and D-204 and a subdomain of the integrins were resolved (PDB IDs 9YBH for M-204, and 9Q68 for D-204)^29^. While our best ranked clusters (cluster 1) did not align well with the crystal structure for M-204, cluster 6 showed good alignment (average DockQ = 0.63^50^) (Supplementary Fig. 4C). However, our modelling procedure successfully predicted the orientation of D-204, with the three best ranked clusters showing an average DockQ score^46^of 0.26, 0.44, and 0.28 (Supplementary Fig. 4D-E). ^47,4827^ For M-204, the VH-Fab glycan site was projected to be outside of the binding pocket, in agreement with our functional data. Importantly, M-204 binds the HPA-1a epitope in a different orientation from D-204 (Fig. 5), which likely explains its capacity to bind two epitopes of HPA-1a and simultaneously crosslink FcγRIIa with the Fc tail causing platelet activation.

**Figure 5.**
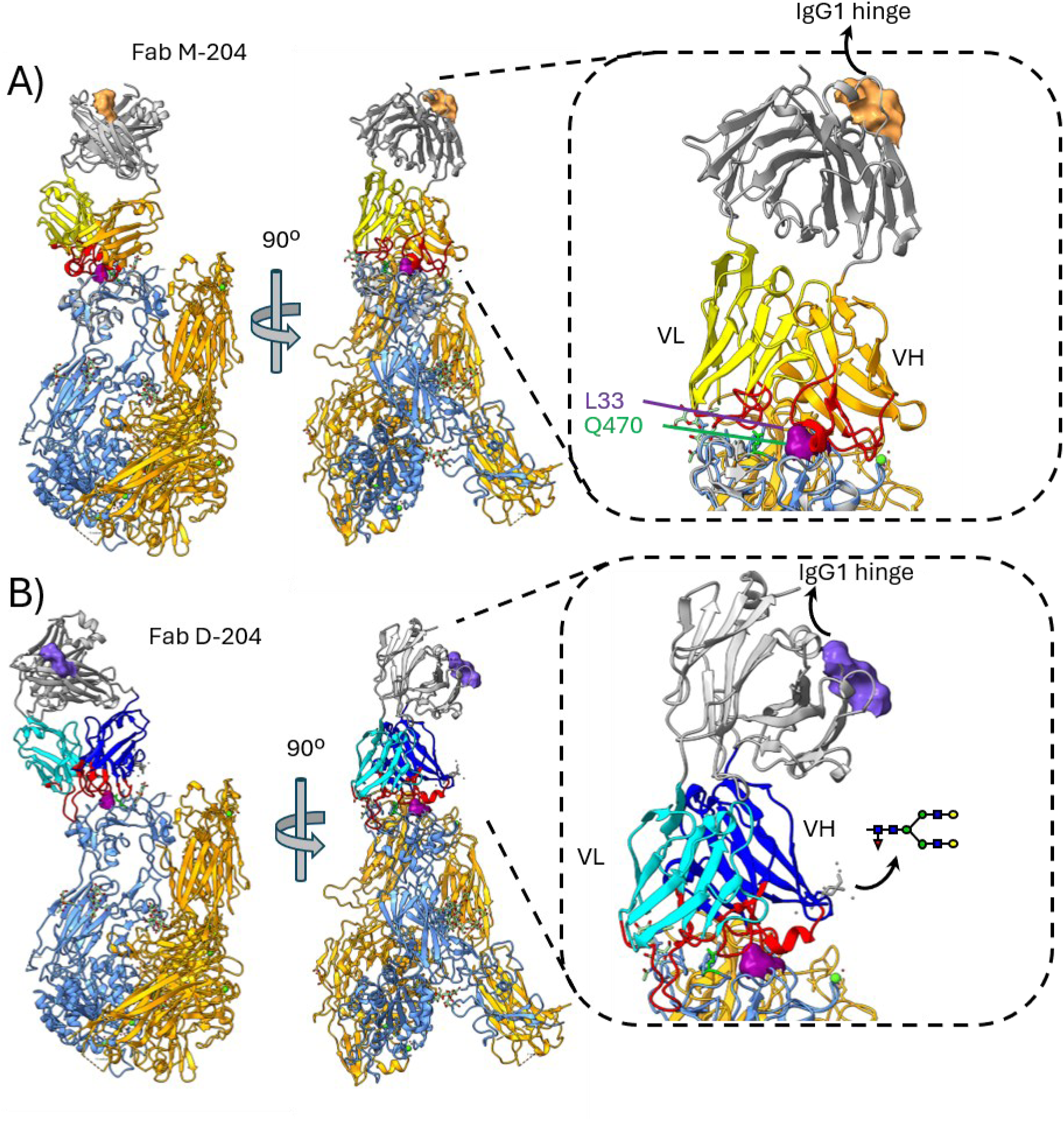
Structural ensembles of M-204 and D-204 docking to integrin αIIbβ3. The crystal miniβ3 structures found in complex with the Fab of **(A)** M-204 (PDB-ID: 9YBH) and **(B)** D-204 (PDB-ID: 9Q68) were aligned to the reported structure of αIIβ3 (PDB-ID: 3FCS). Two views of both structures are shown with 90°C rotation (left and middle panels). An enlarged section highlights the interactions between the HPA-1a epitope and the Fabs, enclosed in a dotted box. The heavy chain residues are colored dark blue or orange, while the light chains are light blue and yellow for M-204 and D-204, respectively. The complementarity regions are marked in red. Annotations include the HPA-1a epitope (L33) indicated by a purple surface rendering; Q470 often contribution to anti-HPA-1a binding is indicated with a green side chain^29^; the VH glycan in M-204. VH: variably heavy, VL: variable light.

## Discussion

There is a growing body of evidence that anti-HPA-1a antibodies, despite targeting the same epitope, can exhibit significant functional heterogeneity^14^. Here, we report the structural and functional characterization of two newly identified anti-HPA-1a antibodies D-and M-204 we identified in recently in a pregnancy characterized by early complications and repeated miscarriages^51^. Our findings reveal distinct characteristics of M-204 which binds to αIIbβ3 reaching saturation earlier compared to other monoclonals. Additionally, M-204 induced platelet aggregation upon binding which was fully inhibited by blocking FcɣRIIa on platelets suggesting that activation occurs through crosslinking of αIIbβ3 with FcɣRIIa. Bivalent binding of both Fab-arms of M-204 was required for this aggregation potential. Perhaps surprisingly, the structural modelling combined with recently available PDB coordinates, the crystal structures suggest similar binding modes of the Fab of M-204 compared to D-204 to HPA-1a. However, they differ in the trajectory where the antibody constant regions would be projected and the second Fab and the Fc, perhaps explaining their differential binding mode with two or one Fab arm, and capacity to engage with FcγRIIa.

Advances in computational molecular modelling have improved structure and interaction prediction significantly in the last years, but do not replace real observations. Our modelling done in parallel with our colleagues who solved the crystal structure^29^ showed that our best scoring structure for D-204 was very close to observations. However, this was not true for M-204, with only the 6^th^ ranked scoring cluster being reflected experimentally. This is a common problem in the antibody-antigen prediction task, as prediction methods are typically able to generate accurate poses but fail to rank them accurately (scoring problem)^52,53^.

Both D-204 and M-204 contain a potential N-glycosylation site in their variable domains, introduced during somatic hypermutation – a feature observed in approximately 15-25% of serum IgG^54^. However, occupation of possible glycan sites is also influenced by adjacent amino acids and overall folding of proteins^55^. The presence of a glycan was confirmed exclusively for M-204. As the most probabilistic models suggest the glycan to be positioned away from the antigen-binding site, it is unlikely to interfere with target binding^29^. This aligns with our data which demonstrates that M-204 engineered without the glycan exhibits identical functional characteristics compared to glycosylated M-204.

The early saturation of M-204 could be explained by several mechanisms. For instance, the membrane organization of platelets and/or conformation of αIIbβ3 could lead to steric hindrance in such a way that binding of M-204 theoretically blocks binding of other M-204 molecules to HPA-1a epitopes in neighboring β3-complexes. A similar phenomenon has also been observed for type II CD20 Fabs which induce steric constraints preventing simultaneous binding of Fabs to CD20 dimers^56^. However, bivalent binding could explain the binding pattern of M-204 to αIIbβ3 which reduces the overall number of antibodies capable of engaging with the target. Bivalent binding has been shown to enhance antibody efficacy in neutralization of SARS-CoV-2, rhinovirus and Dengue neutralization^57^ and is associated with increased complement activation due to target crosslinking^56,58^. M-204 binding kinetics were indeed in agreement with a bivalent interaction mode as binding of a monovalent variant was increased above half-maximum compared to binding seen for all other HPA-1a monoclonals. Monovalent M-204 antibodies also lost their capacity to activate platelets. It is plausible that enhanced target crosslinking could lead to αIIbβ3 activation, outside-in signaling triggering platelet activation, which explains the observed platelet aggregation induced by M-204^12,56^. Furthermore, bivalent binding is likely to confer greater binding stability compared to monovalent interactions, due to lower off-rates, which could amplify signaling^56,59–61^. Curiously, heterozygous HPA-1a/1b platelets, as representative platelets found per definition in alloimmune FNAIT cases, were sensitive to aggregation at a wide range of M-204 concentrations while homozygous HPA-1a/1a platelets were not activated at higher concentrations, an important but not easily explainable observation.

This is the first report of an anti-HPA-1a antibody that is able to directly induce platelet aggregation in an FcɣRIIa-dependent manner. However, a number of studies have reported similar results with other anti-platelet antibodies^62–68^. Notably, an antibody targeting integrin αIIbβ3 was found to cause fibrinogen-mediated aggregation of platelets which was also dependent on an intact Fc tail^68^. In addition, certain anti-HLA antibodies have been shown to activate platelets via FcɣRIIa contributing to decreased platelet survival in refractory patients^62^. Similarly, antibodies targeting glycoprotein Ibα or von Willibrand factor have been demonstrated to induce platelet aggregation which was reversed upon addition of a FcɣRIIa-blocking antibody, suggesting a functional interaction between FcɣRIIa and platelet glycoprotein signaling pathways^63^. Platelet activation through immune complexes and FcɣRIIa engagement has been associated with thrombotic events in patients with severe COVID-19^64^. One could hypothesize that platelet activation by anti-HPA-1a antibodies in FNAIT could lead to decreased platelet survival and risk of thrombotic events in the fetus. Determining whether the characteristics of anti-HPA-1a antibodies in a polyclonal mixture found in patients with M-204-like properties contribute to disease severity in FNAIT remains speculative, as it is unclear how frequently clones arise with properties similar to M-204 in and between patients.

The current data indicate that M-204 preferentially binds αIIbβ3 in a bivalent manner, as suggested by the lower plateau phase observed in the binding curves. We propose that this binding mode enables a specific IgG-Fc orientation that facilitates interaction with nearby FcɣRIIa receptors, either on the same platelet or on adjacent platelets (Fig. 6). Crosslinking of αIIbβ3 and FcɣRIIa activates downstream signaling pathways leading to platelet activation and aggregation. The underlying molecular properties for the functional differences between M-204 and D-204 are not fully understood but is likely influenced by antibody-specific properties such as conformational flexibility and binding orientation^69^. This is likely to also involve active rearrangements as M-204 avidly bound fixed platelets to a similar, or even better than, D-204.

**Figure 6.**
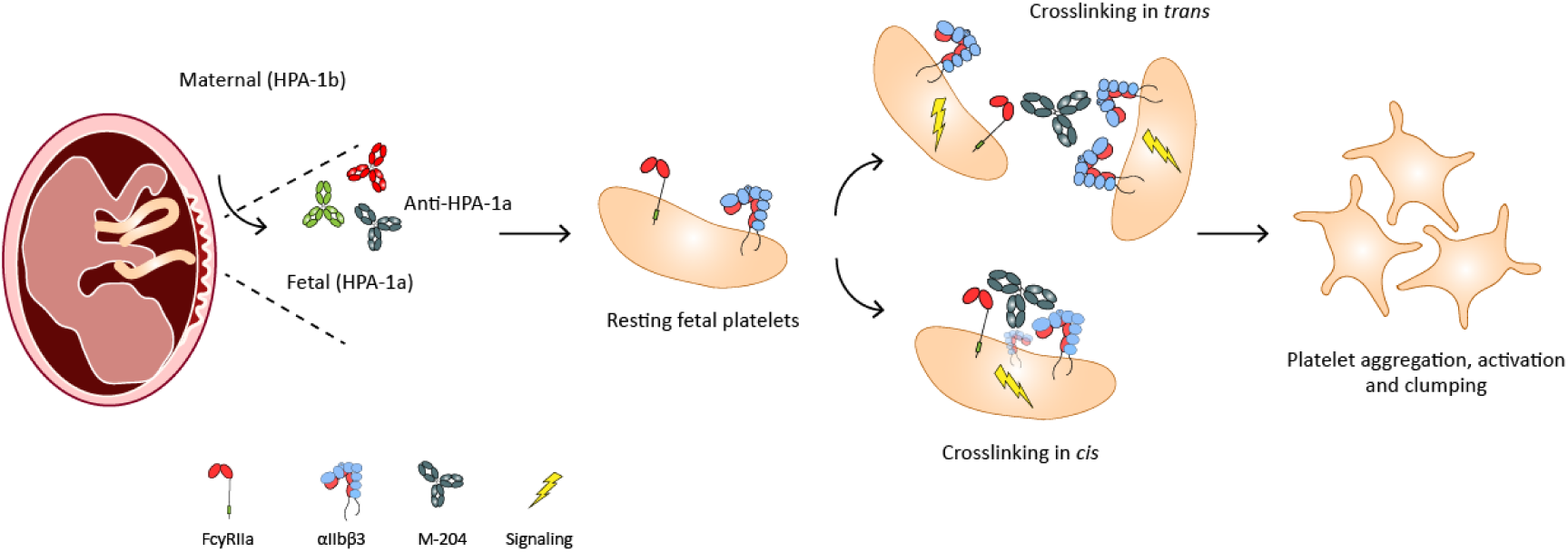
M-204 mediates crosslinking of αIIbβ3 and FcɣRIIa which triggers platelet aggregation. Graphical illustration of the proposed mechanism by which M-204 induces platelet aggregation. M-204 binds the HPA-1a epitope on αIIbβ3 in a bivalent binding manner facilitating interaction of the IgG-Fc with FcɣRIIa in the vicinity. Subsequent crosslinking induces signaling that eventually results in platelet activation and aggregation.

In conclusion, we characterized a novel anti-HPA-1a antibody, M-204, that directly induces platelet aggregation in an FcɣRIIa-dependent manner. M-204 displays distinct structural and functional properties compared to D-204 (and other all currently known HPA-1a specific Mabs), including a Fab glycan, bivalent binding mode to αIIbβ3 and different binding orientation. These characteristics are likely key to its ability to crosslink αIIbβ3 and FcɣRIIa, triggering platelet activation and aggregation. Our findings on monoclonal antibodies reveal functional heterogeneity among anti-HPA-1a antibodies, with M-204 displaying unique integrin-binding and platelet-activating properties. These insights may help to further understand FNAIT pathogenesis.

## Author contribution

JJO, SB, ES and GV designed the study. JJO, SB, MG, RV, IC and CS collected experimental data. JJO wrote the manuscript that was revised by all coauthors. JJO, SB, MG, AB, CS, EDV, SHE, LP, MW, MH, ES and GV contributed to data interpretation and approval of the final version.

## Supplementary figures

**Supplementary figure 1.**
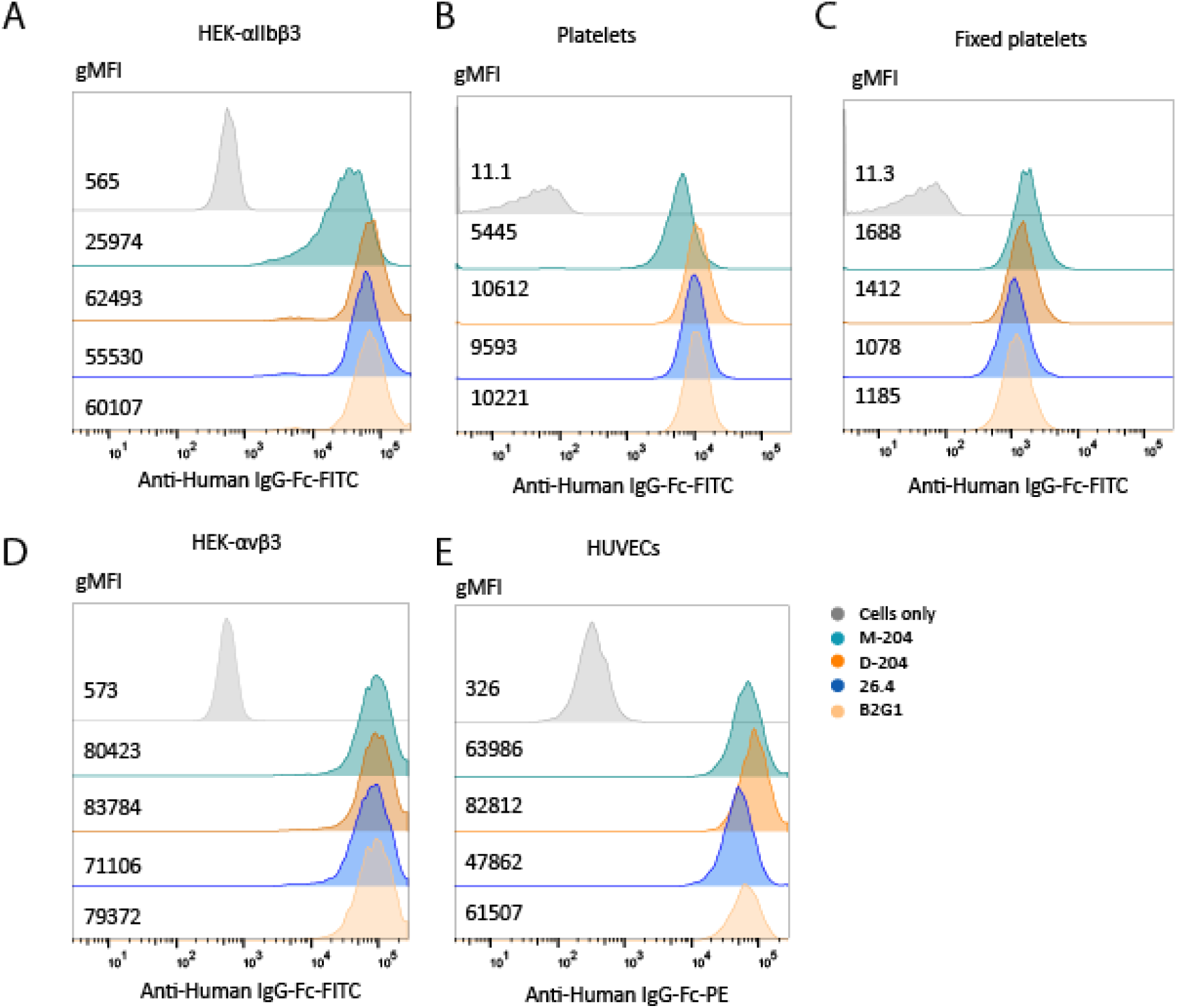
Histograms and absolute gMFIs of monoclonal antibody binding to recombinant HEK-293F cells and cells naturally expressing HPA-1a. Representative histograms comparing raw gMFI values of monoclonal antibody binding to (A) αIIbβ3 on HEK-293F cells, (B) platelets or (C) platelets fixed before staining, and to (D) αvβ3 on HEK-293F cells and (E) HVUECs. The highest concentration of antibodies used in the titrations is shown.

**Supplementary figure 2.**
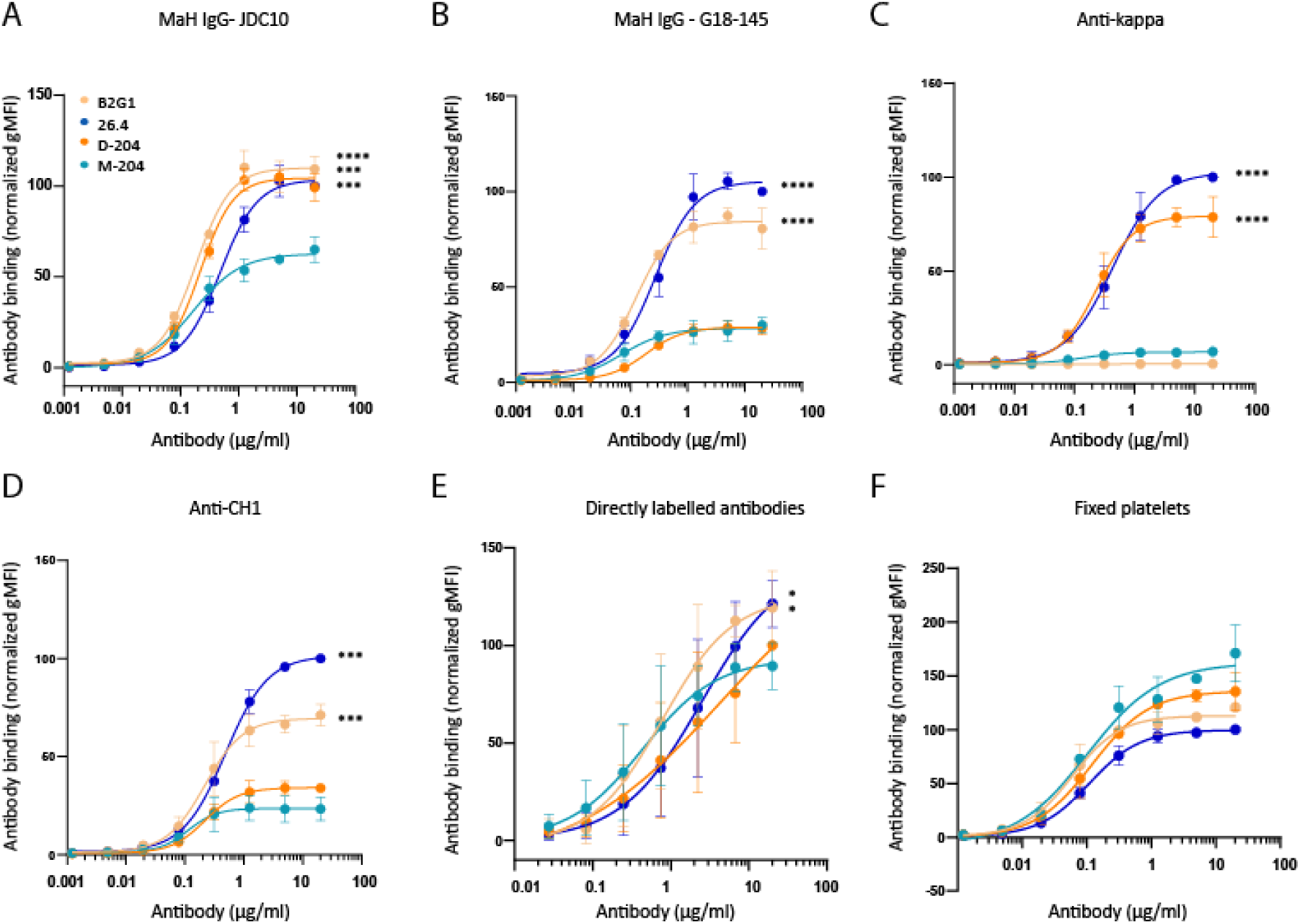
Secondary antibodies and fixation of platelets prior to antibody staining influence saturation plateau of M-204. Different secondary reagents were tested on platelets (HPA-1a/1a) in addition to **(A)** the JDC-10 clone, which was used in the majority of experiments. **(B)** MaH-IgG G18-145 showed a lower saturation plateau for M-204 and D-204. **(C)** The anti-Kappa secondary antibody minimally detected M-204 and no signal was observed for B2G1 due to the presence of Lambda light chain instead of Kappa. **(D)** The anti-CH1 showed a lower saturation for both D-204 and M-204. **(E)** Directly labeled antibodies were titrated on platelets showing a lower plateau phase for M-204. **(F)** platelets were fixed with 1% paraformaldehyde followed by incubation with different monoclonal antibodies detected with JDC10. **(A-F)** Data represents three independent experiments except for **(D)** which is the result of two experiments. Staining of cells was plotted as gMFI normalized towards 20µg/ml 26.4. Error bars indicate standard deviations and statistical analysis was performed on the highest concentration using one-way ANOVA with Dunnet’s multiple comparison test comparing to M-204. Statistically significant differences are indicated by asterisks: ****<0.0001.

**Supplementary figure 3.**
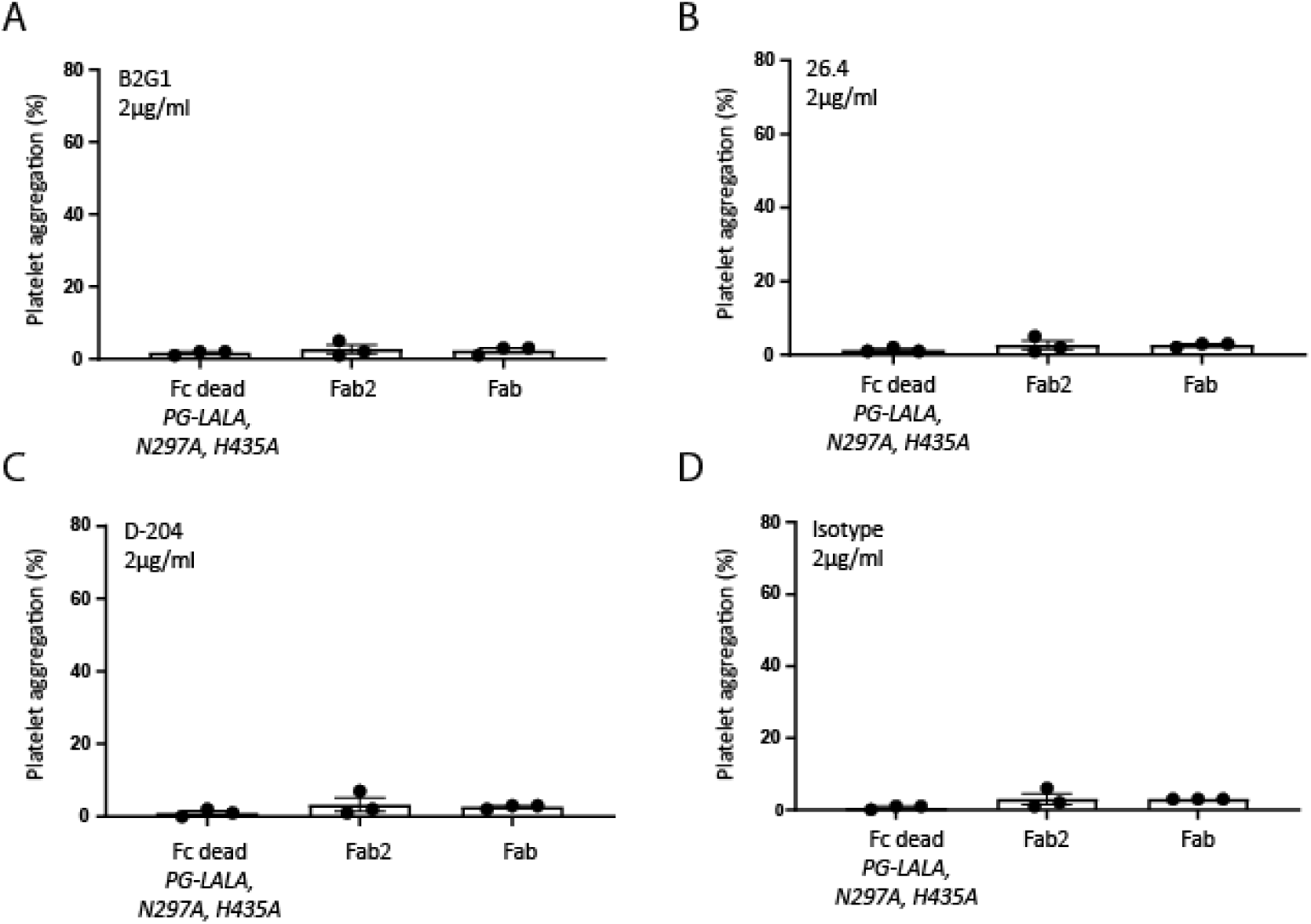
No aggregation of platelets upon addition of different anti-HPA-1a monoclonal antibody formats. (A-D) Different antibody formats of B2G1, 26.4, D-204 and aTNP (isotype) were tested on three different HPA-1a/1a donors for their capacity to induce platelet aggregation. The mean with standard deviation is shown.

**Supplementary figure 4.**
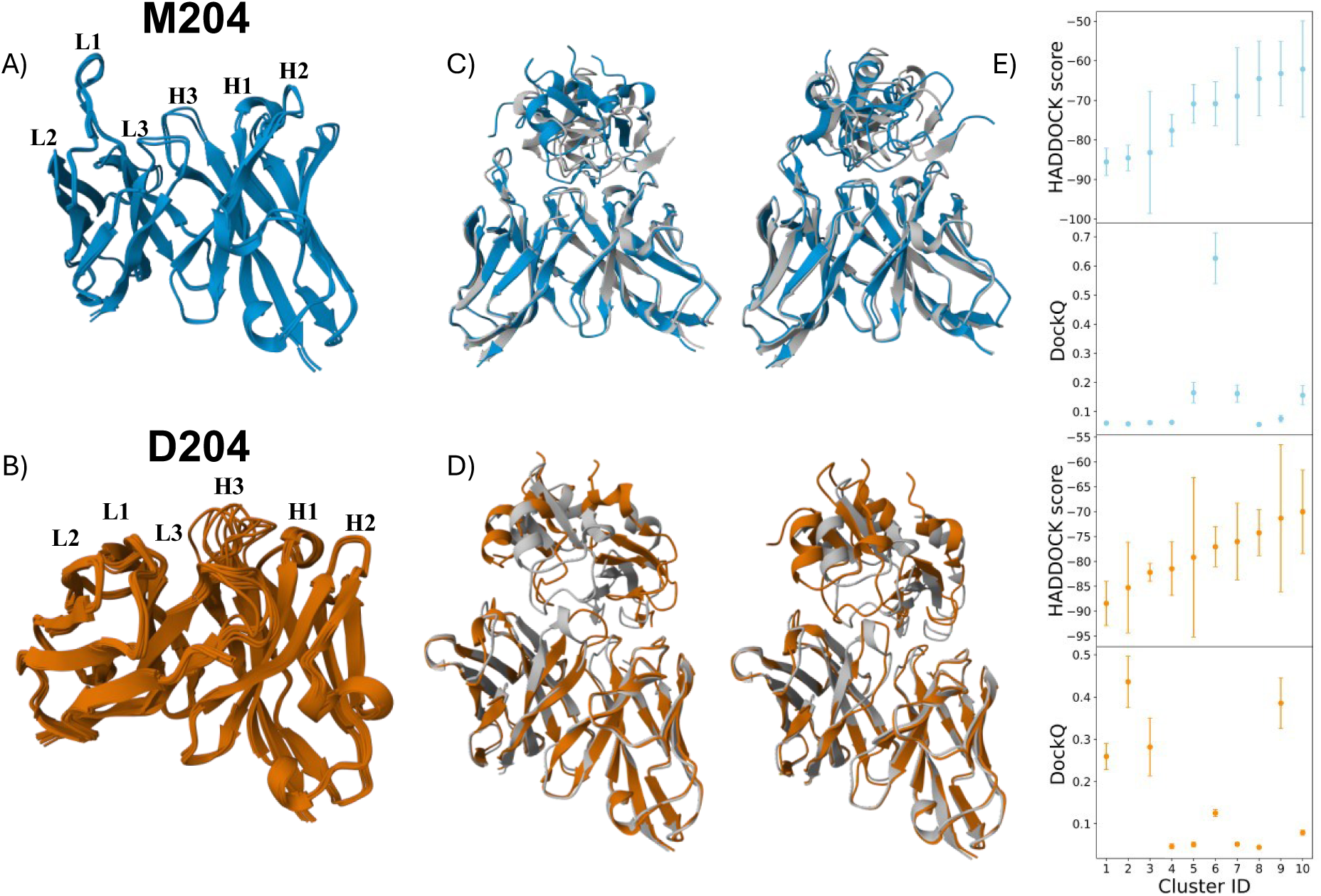
Structural ensembles for M-204 (blue) and D-204 (orange). **(A)** The M-204 ensemble contains only two structures displaying minimal differences across the H3 loop, whereas **(B)** the D-204 ensemble (eight structures) is more heterogeneous, especially on the H3 and L3 loops. **(C)** Comparison between modelled (blue) and crystal structure (PDB-ID:9YBH, gray) for two HADDOCK models of M204. Only the amino acids that are present in the crystal structures are shown, with the peptide backbone encompassing the HPA-1a on top. The first model refers to the best model of the first cluster, while the second model (on the right) depicts the first model of cluster 6, highlighting a substantial agreement with the experimental structure. **(D)** Same comparison but for D204 models (orange) and against the crystal structure (PDB-ID: 9YBH, gray). Here the best ranked models for cluster 1 and 2 are shown. **(E)** HADDOCK and DockQ scores for docking runs using M-204 (blue) and D-204 (orange) for the 10 highest-ranking clusters. The average score and the standard deviations are shown. While the HADDOCK score successfully identifies accurate poses for D204, the most accurate cluster for M204 is ranked sixth.

## References

1. de Vos TW, Winkelhorst D, de Haas M, Lopriore E, Oepkes D. Epidemiology and management of fetal and neonatal alloimmune thrombocytopenia. Transfusion and Apheresis Science. 2020;59(1):102704.

2. Curtis BR. Recent progress in understanding the pathogenesis of fetal and neonatal alloimmune thrombocytopenia. Br J Haematol. 2015;171(5):671–682. doi:10.1111/BJH.13639

3. Kjeldsen-Kragh J, Bengtsson J. Fetal and Neonatal Alloimmune Thrombocytopenia—New Prospects for Fetal Risk Assessment of HPA-1a–Negative Pregnant Women. Transfus Med Rev. 2020;34(4):270–276. doi:10.1016/J.TMRV.2020.09.004

4. Eksteen M, Heide G, Tiller H, et al. Anti-human platelet antigen (HPA)-1a antibodies may affect trophoblast functions crucial for placental development: a laboratory study using an in vitro model. Reprod Biol Endocrinol. 2017;15(1). doi:10.1186/S12958-017-0245-6

5. Yougbaré I, Tai WS, Zdravic D, et al. Activated NK cells cause placental dysfunction and miscarriages in fetal alloimmune thrombocytopenia. Nat Commun. 2017;8(1):224. doi:10.1038/S41467-017-00269-1

6. Bussel JB, Vander Haar EL, Berkowitz RL. New developments in fetal and neonatal alloimmune thrombocytopenia. Am J Obstet Gynecol. 2021;225(2):120–127. doi:10.1016/J.AJOG.2021.04.211

7. Kamphuis MM, Paridaans N, Porcelijn L, et al. Screening in pregnancy for fetal or neonatal alloimmune thrombocytopenia: systematic review. BJOG. 2010;117(11):1335–1343. doi:10.1111/J.1471-0528.2010.02657.X

8. Winkelhorst D, Murphy MF, Greinacher A, et al. Antenatal management in fetal and neonatal alloimmune thrombocytopenia: a systematic review. Blood. 2017;129(11):1538–1547. doi:10.1182/BLOOD-2016-10-739656

9. de Vos TW, Winkelhorst D, Porcelijn L, et al. Natural history of human platelet antigen 1a-alloimmunised pregnancies: a prospective observational cohort study. Lancet Haematol. 2023;10(12):e985–e993. doi:10.1016/S2352-3026(23)00271-5

10. Newman PJ, Derbes RS, Aster RH. The human platelet alloantigens, PlA1 and PlA2, are associated with a leucine33/proline33 amino acid polymorphism in membrane glycoprotein IIIa, and are distinguishable by DNA typing. 1989;83(5). doi:10.1172/JCI114082

11. Huang J, Li X, Shi X, et al. Platelet integrin αIIbβ3: signal transduction, regulation, and its therapeutic targeting. Journal of Hematology & Oncology 2019 12:1. 2019;12(1):1–22. doi:10.1186/S13045-019-0709-6

12. Zhou Y, Wheelock M, Damsky CH, et al. Human cytotrophoblasts adopt a vascular phenotype as they differentiate. A strategy for successful endovascular invasion? 1997;99(9):2139–2151. doi:10.1172/JCI119387

13. Stam W, Wachholz GE, de Pereda JM, Kapur R, van der Schoot E, Margadant C. Fetal and neonatal alloimmune thrombocytopenia: Current pathophysiological insights and perspectives for future diagnostics and treatment. Blood Rev. 2023;59:101038. doi:10.1016/J.BLRE.2022.101038

14. Kroll H, Penke G, Santoso S. Functional heterogeneity of alloantibodies against the human platelet antigen (HPA)-1a. Thromb Haemost. 2005;94(6):1224–1229. doi:10.1160/TH05-03-0159/ID/JR0159-2/BIB

15. Refsum E, Meinke S, Gryfelt G, Wikman A, Höglund P. Adding to the complexity of fetal and neonatal alloimmune thrombocytopenia: Reduced fibrinogen binding in the presence of anti-HPA-1a antibody and hypo-responsive neonatal platelets. Thromb Res. 2018;162:69–76. doi:10.1016/J.THROMRES.2017.12.017

16. Santoso S, Wihadmadyatami H, Bakchoul T, et al. Anti-endothelial αvβ3 antibodies are a major cause of intracranial bleeding in fetal-neonatal alloimmune thrombocytopenia. Arterioscler Thromb Vasc Biol. 2016;36(8):1517. doi:10.1161/ATVBAHA.116.307281

17. Yougbaré I, Lang S, Yang H, et al. Maternal anti-platelet β3 integrins impair angiogenesis and cause intracranial hemorrhage. J Clin Invest. 2015;125(4). doi:10.1172/JCI77820

18. Dardik R, Salomon O. Maternal Anti-HPA-1a Antibodies Increase Endothelial Cell Apoptosis and Permeability. J Vasc Res. 2021;58(5):321–329. doi:10.1159/000515703

19. Thinn AMM, Wang Z, Zhou D, Zhao Y, Curtis BR, Zhu J. Autonomous conformational regulation of β3 integrin and the conformation-dependent property of HPA-1a alloantibodies. Proc Natl Acad Sci U S A. 2018;115(39):E9105–E9114. doi:10.1073/PNAS.1806205115/-/DCSUPPLEMENTAL

20. van Osch TLJ, Oosterhoff JJ, Bentlage AEH, et al. Fc galactosylation of anti-platelet human IgG1 alloantibodies enhances complement activation on platelets. Haematologica. 2022;107(10):2432–2444. doi:10.3324/HAEMATOL.2021.280493

21. de Vos TW, Winkelhorst D, Baelde HJ, et al. Placental Complement Activation in Fetal and Neonatal Alloimmune Thrombocytopenia: An Observational Study. Int J Mol Sci. 2021;22(13). doi:10.3390/IJMS22136763

22. Kapur R, Kustiawan I, Vestrheim A, et al. A prominent lack of IgG1-Fc fucosylation of platelet alloantibodies in pregnancy. Blood. 2014;123(4):471. doi:10.1182/BLOOD-2013-09-527978

23. Sonneveld ME, Natunen S, Sainio S, et al. Glycosylation pattern of anti-platelet IgG is stable during pregnancy and predicts clinical outcome in alloimmune thrombocytopenia. Br J Haematol. 2016;174(2):310–320. doi:10.1111/BJH.14053

24. Stam W, Broekhuis JD, van der Meer FWT, et al. Maternal anti-HPA-1a antibodies block αIIbβ3 and αvβ3 integrin activation which correlates with FNAIT disease severity. Blood. 2026;146(18):2189–2202. doi:10.1182/blood.2025031683

25. Oosterhoff JJ, Bentlage AEH, Falck D, et al. Altered glycosylation profile of anti-HPA-1a-specific antibodies: insights from a prospective fetal and neonatal alloimmune thrombocytopenia cohort. Haematologica. 2026;111(5):1816–1821. doi:10.3324/haematol.2025.288701

26. Oosterhoff JJ, Linty F, Visser R, et al. Generation of human antibodies targeting human platelet antigen (HPA)-1a. Transfusion (Paris). 2024;64(5). doi:10.1111/TRF.17758

27. Zhang H, Paddock C, Oosterhoff JJ, et al. Structural basis of HPA-1a alloimmunization in FNAIT and allosteric regulation of integrin conformation. Blood. 2026;147(22):2569–2581. doi:10.1182/blood.2025032546

28. Eksteen M, Tiller H, Averina M, et al. Characterization of a Human Platelet Antigen-1a–Specific Monoclonal Antibody Derived from a B Cell from a Woman Alloimmunized in Pregnancy. The Journal of Immunology. 2015;194(12):5751–5760. doi:10.4049/JIMMUNOL.1401599

29. Griffin HM, Ouwehand WH. A Human Monoclonal Antibody Specific for the Leucine-33 (P1A1, HPA-1a) Form of Platelet Glycoprotein IIIa From a V Gene Phage Display Library. Blood. 1995;86(12):4430–4436. doi:10.1182/BLOOD.V86.12.4430.BLOODJOURNAL86124430

30. Dekkers G, Treffers L, Plomp R, et al. Decoding the human immunoglobulin G-glycan repertoire reveals a spectrum of Fc-receptor-and complement-mediated-effector activities. Front Immunol. 2017;8(AUG):877. doi:10.3389/fimmu.2017.00877

31. van Osch TLJ, Nouta J, Derksen NIL, et al. Fc galactosylation promotes hexamerization of human IgG1, leading to enhanced classical complement activation. Journal of immunology. 2021;207:1545–1554. doi:10.4049/jimmunol.2100399

32. Labrijn AF, Meesters JI, Priem P, et al. Controlled Fab-arm exchange for the generation of stable bispecific IgG1. Nat Protoc. 2014;9(10):2450–2463. doi:10.1038/NPROT.2014.169

33. Heemskerk N, Gruijs M, Robin Temming A, et al. Augmented antibody-based anticancer therapeutics boost neutrophil cytotoxicity. J Clin Invest. 2021;131(6). doi:10.1172/JCI134680

34. Greenman J, Tutt AL, George AJ, Pulford KA, Stevenson GT, Glennie MJ. Characterization of a new monoclonal anti-Fc gamma RII antibody, AT10, and its incorporation into a bispecific F(ab’)2 derivative for recruitment of cytotoxic effectors. Mol Immunol. 1991;28(11):1243–1254. doi:10.1016/0161-5890(91)90011-8

35. Evans R, O’Neill M, Pritzel A, et al. Protein complex prediction with AlphaFold-Multimer. *bioRxiv*. Published online October 4, 2021:2021.10.04.463034. doi:10.1101/2021.10.04.463034

36. Abramson J, Adler J, Dunger J, et al. Accurate structure prediction of biomolecular interactions with AlphaFold 3. 2024;630(8016):493–500.

37. Abanades B, Wong WK, Boyles F, Georges G, Bujotzek A, Deane CM. ImmuneBuilder: Deep-Learning models for predicting the structures of immune proteins. 2023;6(1):1–8.

38. Dominguez C, Boelens R, Bonvin AMJJ. HADDOCK: A protein-protein docking approach based on biochemical or biophysical information. J Am Chem Soc. 2003;125(7):1731–1737. doi:10.1021/JA026939X/SUPPL_FILE/JA026939XSI20021128_085857.TXT

39. Honorato R V., Trellet ME, Jiménez-García B, et al. The HADDOCK2.4 web server for integrative modeling of biomolecular complexes. 2024;19(11).

40. 3FCS: Structure of complete ectodomain of integrin aIIBb3. Accessed April 16, 2025. https://www.ncbi.nlm.nih.gov/Structure/pdb/3FCS

41. Giulini M, Schneider C, Cutting D, Desai N, Deane CM, Bonvin AMJJ. Towards the accurate modelling of antibody−antigen complexes from sequence using machine learning and information-driven docking. Birol I, ed. Bioinformatics. 2024;40(10):btae583. doi:10.1093/BIOINFORMATICS/BTAE583

42. Raybould MIJ, Turnbull OM, Suter A, Guloglu B, Deane CM. Contextualising the developability risk of antibodies with lambda light chains using enhanced therapeutic antibody profiling. Commun Biol. 2024;7(1). doi:10.1038/S42003-023-05744-8

43. Rodrigues JPGLM, Trellet M, Schmitz C, et al. Clustering biomolecular complexes by residue contacts similarity. Proteins. 2012;80(7):1810–1817. doi:10.1002/PROT.24078

44. Kissel T, Ge C, Hafkenscheid L, et al. Surface Ig variable domain glycosylation affects autoantigen binding and acts as threshold for human autoreactive B cell activation. Sci Adv. 2022;8(6):eabm1759. doi:10.1126/sciadv.abm1759

45. Kaufman EN, Jain RK. Effect of bivalent interaction upon apparent antibody affinity: experimental confirmation of theory using fluorescence photobleaching and implications for antibody binding assays. Cancer Res. Published online 1992.

46. Damelang T, Brinkhaus M, van Osch TLJ, et al. Impact of structural modifications of IgG antibodies on effector functions. Front Immunol. 2024;14:1304365. doi:10.3389/FIMMU.2023.1304365

47. Labrijn AF, Meesters JI, De Goeij BECG, et al. Efficient generation of stable bispecific IgG1 by controlled Fab-arm exchange. doi:10.1073/pnas.1220145110

48. Basu S, Wallner B. DockQ: A Quality Measure for Protein-Protein Docking Models. PLoS One. 2016;11(8):e0161879. doi:10.1371/journal.pone.0161879

49. Oosterhoff JJ, Linty F, Visser R, et al. Generation of human antibodies targeting human platelet antigen (<scp>HPA)</scp>-1a. Transfusion (Paris). 2024;64(5):893–905. doi:10.1111/trf.17758

50. Zhao N, Han B, Zhao C, Xu J, Gong X. ABAG-docking benchmark: a non-redundant structure benchmark dataset for antibody-antigen computational docking. Brief Bioinform. 2024;25(2). doi:10.1093/bib/bbae048

51. Xu X, Coratella I, Reys V, Bonvin AM. DeepRank-Ab: a scoring function for antibody-antigen complexes based on geometric deep learning. Preprint posted online December 6, 2025. doi:10.64898/2025.12.03.691974

52. van de Bovenkamp FS, Hafkenscheid L, Rispens T, Rombouts Y. The Emerging Importance of IgG Fab Glycosylation in Immunity. The Journal of Immunology. 2016;196(4):1435–1441. doi:10.4049/jimmunol.1502136

53. Koers J, Derksen NIL, Ooijevaar-de Heer P, et al. Biased N-Glycosylation Site Distribution and Acquisition across the Antibody V Region during B Cell Maturation. The Journal of Immunology. 2019;202(8):2220–2228. doi:10.4049/JIMMUNOL.1801622

54. Bondza S, Marosan A, Kara S, et al. Complement-Dependent Activity of CD20-Specific IgG Correlates With Bivalent Antigen Binding and C1q Binding Strength. Front Immunol. 2021;11:609941. doi:10.3389/FIMMU.2020.609941/FULL

55. Yan R, Wang R, Ju B, et al. Structural basis for bivalent binding and inhibition of SARS-CoV-2 infection by human potent neutralizing antibodies. Cell Research 2021 31:5. 2021;31(5):517–525. doi:10.1038/s41422-021-00487-9

56. Rougé L, Chiang N, Steffek M, et al. Structure of CD20 in complex with the therapeutic monoclonal antibody rituximab. Science (1979). 2020;367(6483):1224–1230. doi:10.1126/SCIENCE.AAZ9356/SUPPL_FILE/AAZ9356_ROUGE_SM.PDF

57. Teeling JL, French RR, Cragg MS, et al. Characterization of new human CD20 monoclonal antibodies with potent cytolytic activity against non-Hodgkin lymphomas. Blood. 2004;104(6):1793–1800. doi:10.1182/BLOOD-2004-01-0039

58. Bahnan W, Happonen L, Khakzad H, et al. A human monoclonal antibody bivalently binding two different epitopes in streptococcal M protein mediates immune function. EMBO Mol Med. 2023;15(2). doi:10.15252/EMMM.202216208

59. Suzuki A, Yamasaki T, Hasebe R, Horiuchi M. Enhancement of binding avidity by bivalent binding enables PrPSc-specific detection by anti-PrP monoclonal antibody 132. PLoS One. 2019;14(6). doi:10.1371/JOURNAL.PONE.0217944

60. Rijkers M, Saris A, Heidt S, et al. A subset of anti-HLA antibodies induces FcγRIIa-dependent platelet activation. Haematologica. 2018;103(10):1741–1752. doi:10.3324/HAEMATOL.2018.189365

61. Cauwenberghs N, Schlammadinger A, Vauterin S, et al. Fc-receptor Dependent Platelet Aggregation Induced by Monoclonal Antibodies against Platelet Glycoprotein Ib or von Willebrand Factor. Published online 2001.

62. Bye AP, Hoepel W, Mitchell JL, et al. Aberrant glycosylation of anti-SARS-CoV-2 spike IgG is a prothrombotic stimulus for platelets. Blood. 2021;138(16):1481. doi:10.1182/BLOOD.2021011871

63. Nomura Y, Kaneko M, Miyata K, Yatomi Y, Yanagi Y. Bevacizumab and Aflibercept Activate Platelets via FcγRIIa. Invest Ophthalmol Vis Sci. 2015;56(13):8075–8082. doi:10.1167/IOVS.15-17814

64. Mazurov A V., Vinogradov D V., Vlasik TN, Burns GF, Berndt MC. Heterogeneity of Platelet Fc-receptor-dependent Response to Activating Monoclonal Antibodies. Platelets. 1992;3(4):181–188. doi:10.3109/09537109209013181

65. Yu AX, Wu XW, Li JZ, Lian ECY. Mechanism of platelet aggregation induced by a monoclonal antibody requiring Fc portion. Thromb Res. 1993;70(1):51–65. doi:10.1016/0049-3848(93)90223-B

66. Modderman PW, Huisman HG, Van Mourik JA, Von Dem Borne AEGK. A monoclonal antibody to the human platelet glycoprotein IIb/IIIa complex induces platelet activation. Thromb Haemost. 1988;60(1):68–74. doi:10.1055/S-0038-1647637/ID/JR_11/BIB

67. Goldberg BS, Ackerman ME. Underappreciated layers of antibody-mediated immune synapse architecture and dynamics. Yount J, Coelho C, eds. mBio. 2024;16(1). doi:10.1128/MBIO.01900-24

